# Environmental influences on placental programming and offspring outcomes following maternal immune activation

**DOI:** 10.1101/729228

**Authors:** Karen J. Núñez Estevez, Alejandro N. Rondón-Ortiz, Jenny Q.T. Nguyen, Amanda C. Kentner

## Abstract

Adverse experiences during pregnancy induce placental programming, affecting the fetus and its developmental trajectory. However, the influence of ‘positive’ maternal experiences on the placenta and fetus remain unclear. In animal models of early life stress, environmental enrichment (EE) has ameliorated and even prevented associated impairments in brain and behavior. Here, using a maternal immune activation (MIA) model in rats, we test whether EE attenuates maternal, placental and/or fetal responses to an inflammatory challenge, thereby offering a mechanism by which fetal programming may be prevented. Moreover, we evaluate *life-long* EE exposure on offspring development and examine a constellation of genes and epigenetic writers that may protect against MIA challenges. In our model, maternal plasma corticosterone and interleukin-1β were elevated 3 h after MIA, validating the maternal inflammatory response. Evidence for developmental programming was demonstrated by a simultaneous decrease in the placental enzymes *Hsd11b2* and *Hsd11b2/Hsd11b1*, suggesting disturbances in glucocorticoid metabolism. Reductions of *Hsd11b2* in response to challenge is thought to result in excess glucocorticoid exposure to the fetus and altered glucocorticoid receptor expression, increasing susceptibility to behavioral impairments later in life. The placental, but not maternal, glucocorticoid implications of MIA were attenuated by EE. There were also sustained changes in epigenetic writers in both placenta and fetal brain as a consequence of environmental experience and sex. Following MIA, both male and female juvenile animals were impaired in social discrimination ability. Life-long EE mitigated these impairments, in addition to the sex specific MIA associated disruptions in central *Fkbp5* and *Oprm1*. These data provide the first evidence that EE protects placental functioning during stressor exposure, underscoring the importance of addressing maternal health and well-being throughout pregnancy. Future work must evaluate critical periods of EE use to determine if postnatal EE experience is necessary, or if prenatal exposure alone is sufficient to confer protection.

## Introduction

While epidemiological evidence supports an association between maternal infection during pregnancy and risk for central nervous system disorders in offspring (Knuesel et al., 2014), preclinical studies are well poised to inform our mechanistic understanding of these inflammatory mediated outcomes. Indeed, these models have clearly implicated maternal immune activation (MIA) to several neurobiological disruptions that mimic clinical psychiatric pathology (Gumusoglu & Stevens, 2018). For example, MIA is associated with the manifestation of a heterogeneous set of symptoms, including social and cognitive impairments in the offspring later in life, many of which reportedly occur in a sex-dependent manner (CDC, 2014; Kim et al., 2015; Meyer et al., 2014; Reisinger et al., 2015; Zhang et al., 2012).

Although the detrimental effects of prenatal infection are likely due to the maternal, fetal, and/or placental immune and endocrine responses to the infection (Gale et al., 2004; Patterson, 2009; Shi et al., 2005), the specific factors or mechanisms by which the detrimental effects are conferred have yet to be fully elucidated. In rodents, we find that *life-long* exposure to environmental enrichment (EE), a translationally relevant intervention (Woo & Leon, 2013; Woo et al., 2015; Aronoff et al., 2016; Downs et al., 2018; Morgan et al., 2013; Morgan et al., 2015; Purpura et al., 2014), starting prior to breeding and extending through gestation until study’s end protects offspring against some effects of MIA (Connors et al., 2014). However, it is important to determine when during development this intervention is most beneficial and the mechanisms that underlie its positive influence.

An immune challenge during mid-gestation can induce the release of pro-inflammatory immune molecules (i.e. interleukin (IL-1)-1β), the production of stress hormones, and disrupt the expression of placental 11-β hydroxysteroid dehydrogenase (11HSD) 1 and 2, both of which are enzymes critical for glucocorticoid metabolism and the passage of maternal glucocorticoids to the fetus (Diaz et al., 1996; Straley et al., 2014). While 11HSD2 protects the fetus from high maternal levels of glucocorticoids (e.g. cortisol, corticosterone), by rapidly inactivating them into inactive metabolites (e.g. cortisone, 11-dehydrocorticosterone), 11HSD1 converts the inactive glucocorticoids into cortisol/corticosterone (Waddell et al., 1998). Converging basic and clinical evidence suggests that disruptions of these placental enzymes in response to challenge result in excess glucocorticoid exposure to the fetus and altered glucocorticoid receptor expression, increasing susceptibility to behavioral changes later in life (Levitt et al., 1996; Räikkönen et al., 2015; Seckl & Meaney., 2004; Trautman et al., 1995).

One aim of the present study was to determine whether EE exerts its prophylactic effects by attenuating the maternal and/or placental stress and inflammatory responses directly, thereby preventing fetal programming in the offspring. We examined the influence of EE on the release of maternal plasma corticosterone and IL-1β at 3 h (peak of the proinflammatory response) and 24 h post treatment with the inflammatory mediator lipopolysaccharide (LPS) on gestational day (G)15. We also evaluated mRNA expression of proinflammatory and stress-associated molecules in the placenta (e.g. *IL-1β*; *Hsd11b1*, *Hsd11b2*) and fetal brain (e.g. *Nr3c1, Nr3c2*) of male and female offspring. Given that DNA methyltransferase (DNMT)3a and DNA methylation have been implicated in stress-induced downregulation of placental *Hsd11b2* (Jensen Peña et al., 2012), we also evaluated differences in epigenetic machinery recognized to be developmentally important (e.g. DNMT1, DNMT3a, O-GlcNAcylation, methyl cPG binding protein 2; Rose & Klose, 2014; Howerton et al., 2013; Nugent et al., 2018). By evaluating maternal, placental, and fetal responses to prenatal LPS treatment, we aimed to identify the specific level at which EE may offer protection. For example, it is possible that EE may not counteract the maternal inflammatory response but that it instead a) prevents the release of placental cytokines such as IL-1β, or b) contributes to the maintenance of glucocorticoid metabolism via placental *Hsd11b1* and *Hsd11b2*. If so, this would be the first report that EE could specifically protect placental function at the time of a maternal stressor, without affecting maternal responses to stress.

A second aim was to extend upon our previous findings that life-long EE can attenuate MIA induced social impairments in male offspring. While male, but not female, rats demonstrated impairments in the social interaction test (Connors et al., 2015), we hypothesized that the apparent resiliency of MIA exposed females may be dependent on the behavioral and central endpoints evaluated; indeed, males and females may present with differing phenotypes following early life challenges (Goldstein et al., 2019). We also tested whether exposure to complex housing environments could be beneficial for females following MIA.

## Methods

### Animals and housing

Sprague Dawley rats were acquired from Charles River Laboratories (Wilmington, MA), and housed at 20°C on a 12 h light/dark cycle (0700-1900 light) with *ad libitum* access to food and water. A schematic timeline of experimental procedures is presented in **Figure 1A.**Female rats were housed in pairs in one of two conditions: environmental enrichment (EE; 91.5 × 64 × 159 cm; see **Figure 1B**), comprised of large multi-level cages with ramps and access to toys, tubes, chew bones, and Nestlets® (Ancare, Bellmore, NY), or standard cages (SD; 27 × 48 × 20 cm; **see Figure 1C**). Male rats were paired in SD conditions unless they were breeding, at which point they were housed with two females in EE or larger SD one level cages (51 × 41 × 22 cm) with access to one tube, one chew bone and Nestlets® (Ancare, Bellmore, NY). For the EE cages, toys, tubes and chew bones were switched out twice weekly. The current EE regimen was employed as we have previously demonstrated that it has utility in both the protection (Connors et al., 2014) and reversal (Kentner et al., 2016) of LPS-induced MIA effects. In accordance to recent guideline recommendations for improving the reporting of MIA model methods, we have completed the reporting table from Kentner et al. (2019a) and provided it as ***Supplementary Table 1***. Animal procedures were approved by the MCPHS University Institutional Animal Care and Use Committee and carried out in compliance with the recommendations outlined by the Guide for the Care and Use of Laboratory Animals of the National Institutes of Health.

**Table 1.** Central mRNA gene expression of fetal epigenetic writers (juvenile time point).

| Sex | Housing | Gestational Treatment | Hippocampus |  |  |  | Hypothalamus |  |  |  |
| --- | --- | --- | --- | --- | --- | --- | --- | --- | --- | --- |
|  |  |  | *OGT* | *DNMT1* | *DNMT3a* | *MECP2* | *OGT* | *DNMT1* | *DNMT3a* | *MECP2* |
| Male | SD | Saline | 1.00±0.12 | 1.00±0.06 | 1.00±0.07 | 1.00±0.05 | 1.00±0.15 | 1.00±0.08 | 1.00±0.08 | 1.00±0.04 |
|  |  | LPS | 0.99±0.22 | 0.84±0.08 | 0.85±0.07 | 0.96±0.10 | 1.07±0.13<sup>aa</sup> | 1.09±0.13<sup>aaa</sup> | 0.99±0.10<sup>aaaa</sup> | 1.22±0.14 |
|  | EE | Saline | 0.84±0.15 | 0.83±0.20 | 0.87±0.18 | 0.92±0.14 | 1.39±0.16 | 1.15±0.07 | 0.86±0.04 | 1.50±0.23<sup>bbbb</sup> |
|  |  | LPS | 0.59±0.07 | 0.65±0.05 | 0.89±0.05 | 0.96±0.13 | 0.90±0.14<sup>aa</sup> | 0.83±0.7<sup>aaa</sup> | 0.63±0.02<sup>aaaa</sup> | 0.93±0.05<sup>bbbb</sup> |
| Female | SD | Saline | 1.00±0.06 | 1.00±0.06 | 1.00±0.11 | 1.00±0.05 | 1.00±0.12 | 1.00±0.12 | 1.00±0.066 | 1.00±0.03<sup>bbbb</sup> |
|  |  | LPS | 0.94±0.09 | 0.97±0.18 | 0.94±0.11 | 0.97±0.11 | 1.09±0.15<sup>aa</sup> | 0.66±0.04<sup>aaa</sup> | 0.76±0.05<sup>aaaa</sup> | 0.83±0.06<sup>a, bbbb</sup> |
|  | EE | Saline | 1.15±0.18 | 1.10±0.20 | 1.11±0.17 | 1.01±0.09 | 4.86±0.39 | 1.14±0.05 | 1.19±0.05 | 5.75±0.95 |
|  |  | LPS | 1.31±0.12 | 1.06±0.10 | 0.98±0.06 | 0.96±0.08 | 0.83±0.14<sup>aa</sup> | 0.60±0.04<sup>aaa</sup> | 0.58±0.05<sup>aaaa</sup> | 0.68±0.04<sup>a</sup> |

Error bars represent mean ± SEM, n = 7-8 per group. <sup>a</sup>p < 0.05, <sup>aa</sup>p < 0.01, <sup>aaa</sup>p < 0.001, <sup>aaaa</sup>p < 0.0001, significantly different from saline; <sup>b</sup>p < 0.05, <sup>bb</sup>p < 0.01, <sup>bbb</sup>p < 0.001, <sup>bbbb</sup>p < 0.0001 significantly different from female EE. Data normalized to same-sex SD-Saline groups.

**Figure 1.**
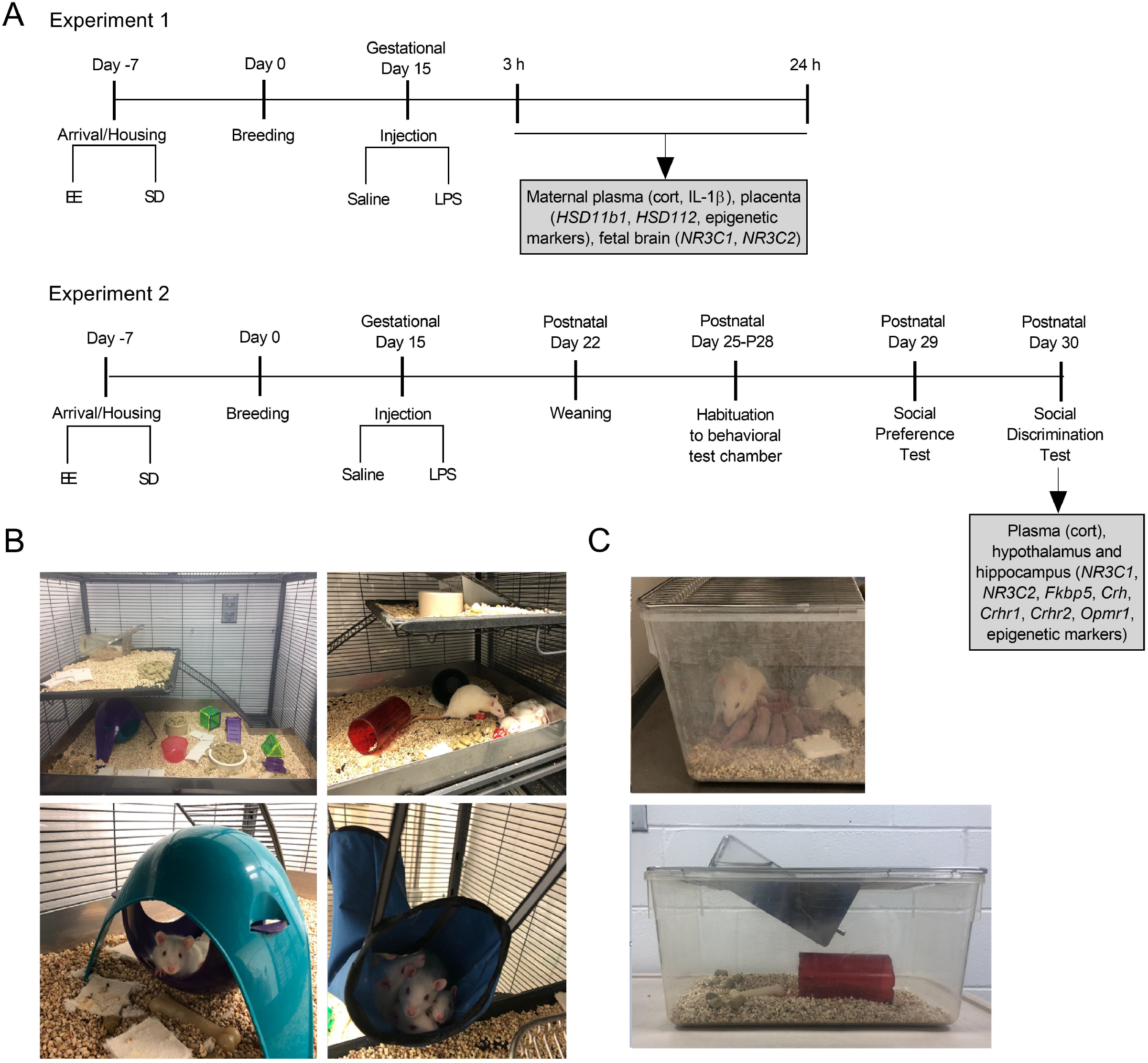
A) Schematic representation of timeline and experimental design. **Experiment 1**. Female rats habituated to either standard (SD) or environmentally enriched (EE) housing and bred. Pregnant rats received an injection of LPS (100 μg/kg) or equivolume of saline on gestational day 15. 3 h and 24 h after injection, plasma and tissue samples were collected from placenta, fetus and pregnant rats to analyze corticosterone, *Hsd11b1*, *Hsd11b2*, *Nr3c1*, *Nr3c2*, and epigenetic writers. **Experiment 2**. Pregnant SD and EE rats received an injection of LPS (100 μg/kg) or equivolume of saline on gestational Day 15. After birth on postnatal day (P)22, offspring were weaned. Between P25-P28, offspring were habituated to an open arena where social behavior tests would take place. On P29 and P30 the social preference and social discrimination tests were conducted, respectively. On the afternoon following behavioral testing on P29, offspring plasma, hippocampus and hypothalamus samples were collected for future analyses. B) Example set-ups of the multi-level environmental enrichment caging system. C) Representative photographs of the standard cages used in the present study.

### Breeding and validation of maternal immune activation (MIA)

Following a week-long acclimatization to their housing conditions, female rats were bred and pregnancy was confirmed by the presence of spermatozoa in vaginal samples combined with a sustained period of diestrus and constant weight gain. On gestational day (G)15, dams received an i.p. injection of 100 μg/kg of LPS (*Escherichia coli* serotype 026:B6;L-3755, Sigma, St. Louis, MO) or an equivalent volume of pyrogen free saline. The choice of rat strain, dose and timing of LPS administration was based on findings that these conditions result in a robust inflammatory response, as demonstrated by significant weight loss and sickness behaviors (Connors et al., 2015; Kentner et al., 2016). Two groups of dams (N = 56; **Figure 1A, Experiment 1**) were euthanized by ketamine/xylazine (40–80 mg/kg, i.p./5–10 mg/kg, i.p.) at either 3 h or 24 hr following the G15 challenge. Maternal plasma samples were collected and stored for later validation of MIA by corticosterone and interleukin (IL)-1β levels. Dams were perfused intracardially with a phosphate buffer solution (PBS) and the uterine horn removed. Placentas (as previously described by Jensen Peña et al., 2012), whole fetal brains, and bodies were saved for DNA and RNA isolation and sex identification. To limit effects of intrauterine position, only the first three conceptuses from either side of the cervical end were selected (Bronson & Bale, 2014).

A third set of dams (N= 37), also treated with LPS or saline (**Figure 1A, Experiment 2**), were permitted to continue their pregnancy through until birth (postnatal day (P)1). On G19, SD dams were separated into individual cages (27 × 48 × 20 cm; **see Figure 1C**), while a physical divider separated EE dams (allowing for olfactory, tactile, auditory, and some visual contact; important components of EE), for the purpose of preventing the mixing of litters. On P2, pups were culled to 12 pups per litter (6 males and 6 females, wherever possible). Offspring were maintained in their respective housing conditions with their mothers until weaning on P22. At that point, offspring were placed into clean cages, maintaining their same neonatal housing conditions until behavioral testing.

### Behavioral Tests

Between P25-P28, one male and one female offspring from each litter were habituated daily in an open field arena (40 cm × 40 cm ×28 cm) which was used to evaluate social behaviors on P29-P30. This developmental time point was selected as the frequency of social behaviors peak during the juvenile and early adolescent period (Thor & Holloway, 1984). All equipment was thoroughly cleaned with Quatriside TB between each animal. For all behavioral tests, videos were scored via an automated behavioral monitoring software program (Cleversys TopScan, Reston, VA) and evaluated for total distance (mm) and velocity (mm/sec) traveled in the arena.

#### Social Preference

Using the TopScan automated scoring software, the arena was delineated into three equal-sized ‘zones’. Juvenile P29 rats were placed into the center of the arena (zone 1) and evaluated on their choice to visit a novel inanimate object (zone 2) or an unfamiliar same sex/size/age conspecific (zone 3), each contained within a small wire cup on opposite ends of the arena (Trial 1; Crawley, 2007). The frequency of crosses into each of the zones and the duration spent visiting was automatically recorded by the TopsScan software.

#### Social discrimination

24 h later, rats were returned to the choice task, this time one chamber contained the same unfamiliar conspecific from the day before, and the other chamber held a novel animal of the same sex/size/age (Trial 2). For both Trials 1 and 2, the following parameters were analyzed: total investigation time of both chambers (s), total time in *social investigation* (s), and either a social preference or discrimination ratio (duration of time (s) with novel rat / duration of time (s) with object (Trial 1 – *social preference*) or familiar rat (Trial 2 – *social discrimination*). Higher scores indicated more time spent with the novel rat.

### P30 Offspring Tissue collection

Following the social discrimination test, P30 offspring were anesthetized with a mixture of ketamine/xylazine (40–80 mg/kg, i.p. / 5–10 mg/kg, i.p.). Cardiac blood was collected and stored frozen as plasma. Animals were perfused intracardially with PBS, brains removed, placed over ice and the hippocampus and hypothalamus dissected. All samples were immediately frozen on dry ice and stored at −80 °C until analysis.

### ELISA

ELISA was used to analyze plasma corticosterone levels using the small sample assay protocol provided with the manufacturer instructions (ADI-900-097, Enzo Life Sciences, Farmingdale, NY, USA). The minimum detectable concentration was 26.99 pg/ml, and the intra and inter-assay coefficients of variation were 6.6% and 7.8%, respectively. The minimum detectable dose of rat IL-1β (R&D Systems) was 5 pg/mL, and the intra and inter-assay coefficients of variation ranged between 3.9-8.8% and 4.0-5.7%, respectively. All ELISA samples were run in duplicate.

### Extraction of genomic DNA and polymerase chain reaction

Genomic DNA was extracted from embryonic tissue (e.g. fetal body) using a DNA extraction kit (DNeasy Blood & Tissue Kits, QIAGEN, Valencia, CA, USA), according to manufacturer’s instructions. Polymerase chain reaction (PCR) was done using 1μL of genomic DNA and *Sry* primers to yield a 317 base pair product. The sequence of the *Sry* forward primer was: 5’CGGGATCCATGTCAAGCGCCCCATGAATGCATTTATG 3’ and reverse primer 5’ GCGGAATTCACTTTAGCCCTCCGATGAGGCTGATAT 3’. As a control the sequence of beta actin (ACTB, Chr. 12p11) forward primer 5’ AGCCAT GTACGTAGCCATCC 3’ and reverse primer 5’ TGTGGTGGTGAAGCTGTAGC 3’ was amplified by PCR to yield a 240-bp product (Levine et al., 2000). The PCR was done by adding 22.6 μL of DNA PCR Supermix (Promega, Madison, WI, USA) to a PCR tube, then adding 0.5μL of SRY forward primer and 0.5 μL of SRY reverse primer. 0.5 μL of ACTB forward primer (1:10 dilution) and 0.5 μL of ACTB reverse primer (1:10 dilution) was added to the PCR tube along with 1μL of DNA. The PCR conditions were preheating at 105°C, then setting initial denature to 95 °C for 2 min. The PCR machine was set to 35 cycles, step 1: 95°C (1 min), step 2: 52°C (1 min) step 3: 72°C (1 min), final extension: 72°C (5 min) and a final hold at 10°C. The product was loaded onto a 1.8% agarose gel and sex was determined by presence of *Sry* band (**Figure 2A**; adapted from Miyajima et al., 2009).

**Figure 2.**
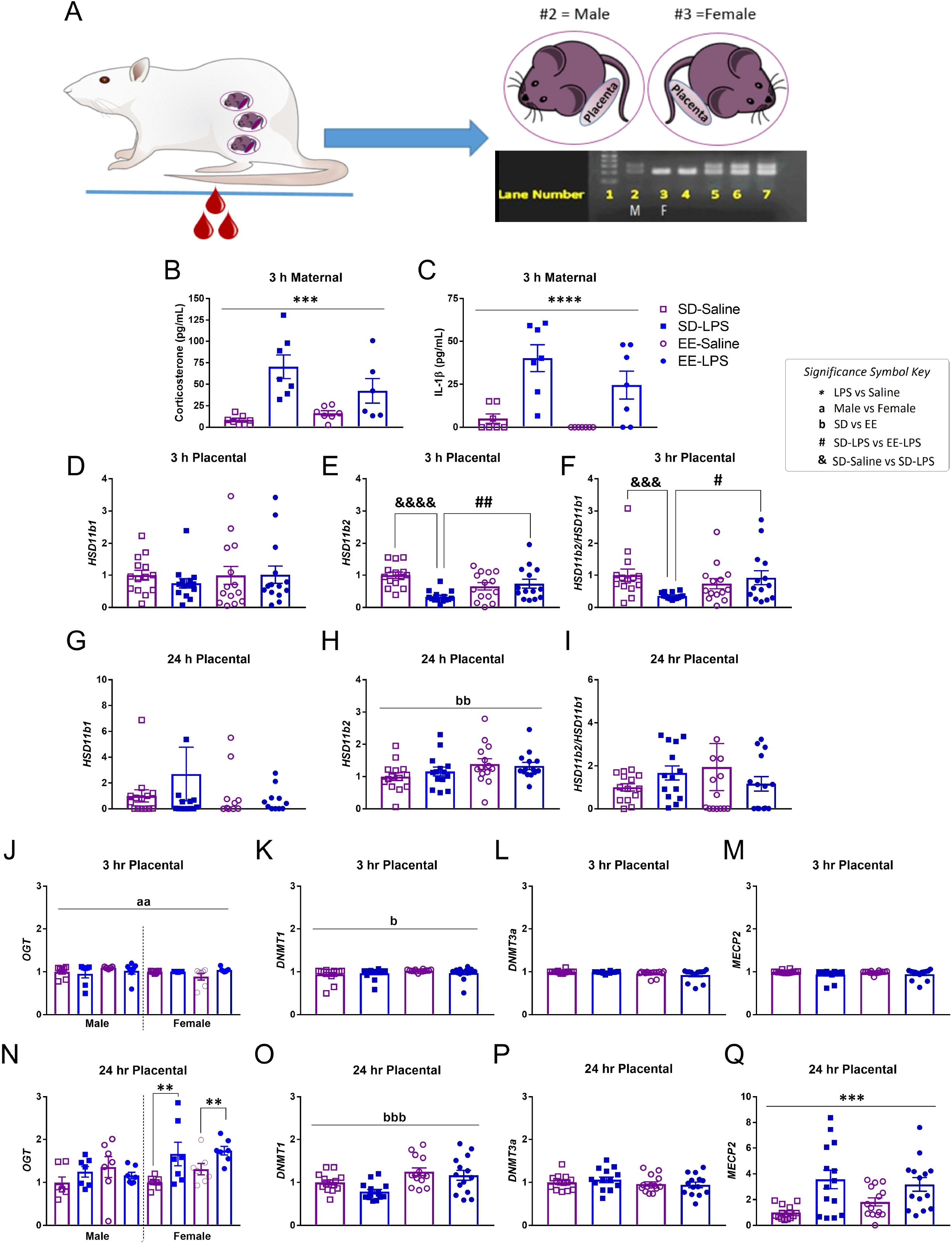
A) Illustration of maternal blood collection, embryonic tissue collection and PCR to determine fetal sex. Lane (L) 1; molecular weight ladder. L2 is the ‘known’ male (M) control sample, confirmed by the presence of the top band (*Sry* gene). L3 is the ‘known’ female (F) control, as depicted by an absence of *Sry*. L4 = female; L5-7 = males. Maternal plasma levels of B) corticosterone (pg/mL) and C) IL-1β (pg/mL) were elevated 3 h post LPS, validating maternal immune activation. Placental levels of D,G) *Hsd11b1*, E,H) *Hsd11b2*, F,I) *Hsd11b2*/*Hsd11b1*, J,N) *OGT*, K,O) *DNMT1*, L,P) *DNMT3a*, M,Q) *MECP2*, 3 h and 24 h post LPS. Group designations: SD-Saline (open squares), SD-LPS (closed squares), EE-Saline (open circles), EE-LPS (closed circles). Error bars represent mean ± SEM, n = 7 per sex/housing/MIA group. If there were no significant effects of sex, male and female offspring data were collapsed together for visualization. *p < 0.05, ** p < 0.01, *** p < 0.001, ****p <0.0001, LPS vs saline; ^a^p < 0.05, ^aa^p < 0.01, ^aaa^p < 0.001, ^aaaa^p <0.0001, male vs female; ^b^p < 0.05, ^bb^p < 0.01, ^bbb^p < 0.001, ^bbbb^p <0.0001, standard housing vs environmental enrichment; ; ^&^p < 0.05, ^&&^p < 0.01, ^&&&^p < 0.001, ^&&&&^p <0.0001, SD-Saline vs SD-LPS; ^#^p < 0.05, ^##^p < 0.01, SD-LPS vs EE-LPS. Fetal brain data are presented in ****Supplementary Table 2****.

### RNA Isolation and Gene Expression Analysis

For fetal and juvenile RNA isolation, tissue was added to TRIzol Reagent (Invitrogen) and briefly homogenized before proceeding with chloroform precipitation and isopropanol. After the RNA pellet wash with ethanol, RNA was re-suspended in 50 μL of RNase-free water and concentration of RNA was determined by absorbance at 260 nm using (ND-1000 Spectrophotometer, NanoDrop Technologies, Wilmington, DE, USA).

The RNA (150 ng) was transcribed to complementary DNA (cDNA) using the Transcriptor First Strand cDNA Synthesis Kit (Roche, Risch-Rotkreuz, Switzerland). Gene expression changes in IL-1β (Rn00580432_m1), glucocorticoid receptor (*Nr3c1*; Rn00561369_m1), FK506 binding protein 5 (*Fkbp5*; Rn01768371_m1), mineralcorticoid receptor (*Nr3c2*; Rn00565562_m1), 11-beta hydroxysteroid dehydrogenase type 1 (*Hsd11b1*; Rn00567167_m1), 11-beta hydroxysteroid dehydrogenase type 2 (*Hsd11b2*; Rn00492539_m1), corticotrophin releasing hormone (*Crh*; Rn01462137_m1), corticotrophin releasing hormone receptor type 1 (*Crhr1*; Rn00578611_m1), corticotrophin releasing hormone receptor type 2 (*Crhr2*; Rn00575617_m1), μ-opioid receptor (*Oprm1*; Rn01430371_m1), DNA methyltransferase 1 (*DNMT1*; Rn00709664_m1), DNA methyltransferase 3 alpha (*DNMT3a*; Rn01027162_g1), methyl cPG binding protein 2 (*MECP2*; Rn01529606_g1), and O-GlcNAcylation (*OGT*; Rn00820779_m1) were assessed using the TaqMan Gene Expression Assay (Applied Biosystems, Norwalk, CT). Glyceralaldehyde-3-phosphate dehydrogenase (*Gapdh*; Rn99999916_s1) was used as an endogenous control for each sample. All of the samples and controls were done in triplicate wells in 96-well plates and were analyzed using the Applied Biosystems 7500 Fast Real-Time PCR (Norwalk, CT). The standard curve method was utilized as previously described (Kentner et al., 2018a), using semi quantitative analysis of gene expression normalized to *Gapdh*. Data are presented at mean expression relative to same-sex SD-Saline treated rats.

### Statistical analyses

Statistics were performed using the software package Statistical Software for the Social Sciences (SPSS) version 21.0 (IBM, Armonk, NY). Data points that fell either above or below 2 standard deviations of the mean were removed (no more than one value per group; Miller, 1991; Howland et al., 2012). Alpha levels were set to p< 0.05 for all omnibus tests. One (gestational treatment × housing) and three-way (sex × gestational treatment × housing) ANOVAs were conducted, as appropriate. When data were significantly skewed (Shapiro-Wilks), Kruskal-Wallis tests (non-parametric equivalent to ANOVA expressed as *Χ^2^*) were employed, followed by Mann-Whitney U tests (expressed as *z;* see Green & Salkind, 2005). Pairwise comparisons were made using conservative adjusted alpha correction values. With respect to parametric data, LSD post hocs were applied except where there were fewer than three levels, in which case pairwise t-tests and Levene’s (applied in the occurrence of unequal variances on the post hoc assessments) were utilized. The False Discovery Rate (FDR) was applied to correct for the multiple testing in gene expression experiments. Male and female offspring data were collapsed together, unless there was a significant sex difference, and data graphically expressed as mean ± SEM.

## Results

### Validation of MIA

There was a significant effect of gestational treatment on maternal plasma corticosterone (F(1, 23) = 20.420, p = 0.0001; **Figure 2AB**) and IL-1β (*X*^2^(1) = 13.906, p =0.0001; **Figure 2AC**) at 3 h, but not 24 h (p >0.05; data not shown) following LPS challenge, validating both inflammation and high circulating levels of maternal glucocorticoids. Moreover, in the third cohort of dams (those permitted to give birth), LPS treated animals had significantly slowed body weight gain (−0.37 ± 1.02 g) between G15 and G16 compared to saline (5.07 ± 1.65 g; F(1, 33) = 5.767, p = 0.022).

### MIA and enrichment effect placental enzymes and epigenetic writers

Three hours following MIA, male and female placental *Hsd11b1* were intact (p >0.05; **Figure 2D**). However, *Hsd11b2* (*X*^2^(1) = 7.047, p = 0.008; **Figure 2E**) and *Hsd11b2*/*Hsd11b1* (a general indicator for placental glucocorticoid turnover, Heussner et al., 2016; *X*^2^(1) = 5.114, p = 0.024; **Figure 2F**) were both significantly diminished compared to saline (p = 0.001) in placenta of both sexes. Given that the major purpose of our study was to evaluate the potential for EE to protect against the effects of MIA, we conducted an *a priori* test and demonstrated that complex housing attenuated MIA mediated effects on placental *Hsd11b2*. Specifically, placenta *Hsd11b2* (*z* = −4.181, p = 0.0001; **Figure 2E)**and *Hsd11b2/Hsd11b1* (*z* = −3.170, p = 0.002; **Figure 2F)**were significantly lower in SD-LPS treated animals compared to SD-Saline. Placenta *Hsd11b2* (*z* = −2.435, p = 0.014) and *Hsd11b2/Hsd11b1* (*z* = −2.160, p =0.031) were also significantly lower in SD-LPS rats compared to EE-LPS. There were no significant differences between EE-Saline and EE-LPS treated rats (p >0.05) on either of these measures.

Twenty-four hours later, while placental *Hsd11b1* (**Figure 2G**), *Hsd11b2* (**Figure 2H**) and *Hsd11b2*/ *Hsd11b1* (**Figure 2I**) were normalized as a function of gestational treatment, *Hsd11b2* levels were significantly elevated following EE (*X*^2^(1) = 6.286, p = 0.012; **Figure 2H**).

Three hours following MIA, female placentas had significantly higher levels of *OGT* than males (*X*^2^(1) = 6.619, p = 0.010; females: 1.22 +/−0.03 and males: 1.15 +/− 0.04; **Figure 2J**). Enrichment was associated with elevated placental *DNMT1* (*X*^2^(1) = 5.491, p =0.019; EE: 1.15 +/− 0.02 and SD: 1.10 +/− 0.03; **Figure 2K**), however *DNMT3a* (**Figure 2L**) and *MECP2* (**Figure 2M**) were unaffected by early life experience at this time point (p>0.05).

A significant interaction of sex by gestational treatment was revealed for placental *OGT* 24 h following MIA (F(1, 48) = 4.441, p = 0.040; **Figure 2N**). Female LPS treated placentas had elevated *OGT* compared to their saline treated counterparts (t(26) = 3.368, p = 0.002). This is suggestive of a compensatory protective effect in females following early life adversity. Placental *DNMT1* (F(1, 48) = 15.856, p = 0.0001;; **Figure 2O**) was higher in EE animals compared to SD 24 h post MIA (t(54) = −3.805, p = 0.0001). *DNMT3a* was unaffected (p>0.05; **Figure 2P**) while placental *MECP2* was increased 24 h following MIA (*X*^2^(1) = 14.082, p = 0.0001; **Figure 2Q**).

### Enrichment effects on GR mRNA in the fetal brain

There were no significant effects on either fetal *Nr3c1* or *Nr3c2* (p >0.05) 3 h post MIA. However, 24 h later, EE animals had elevated levels of *Nr3c1* (*X*^2^(1) = 12.413, p = 0.0001) and *Nr3c2* (*X*^2^(1) = 17.598, p = 0.0001) compared to SD rats (***Supplementary Table 2***).

### MIA and enrichment effect juvenile behavior

There were main effects of housing and gestational treatment on distance traveled (**Figure 3AB**) and velocity (**Figure 3C**) during Trial 1 of the juvenile social behavioral tests (see **Supplementary Table 3**). The reduced travel (p = 0.001) and speeds (p = 0.004) exhibited by the EE group were not unexpected as this rearing condition is classically associated with quick habituation to novel environments (Kentner et al., 2018a). LPS treated animals traveled at greater speeds (p = 0.001) and distances (p = 0.028) compared to saline, suggestive of a hyperactive phenotype. Duration of time spent in the center of the social test chamber was unaffected by early life experience (p>0.05; **Figure 3D**).

**Figure 3.**
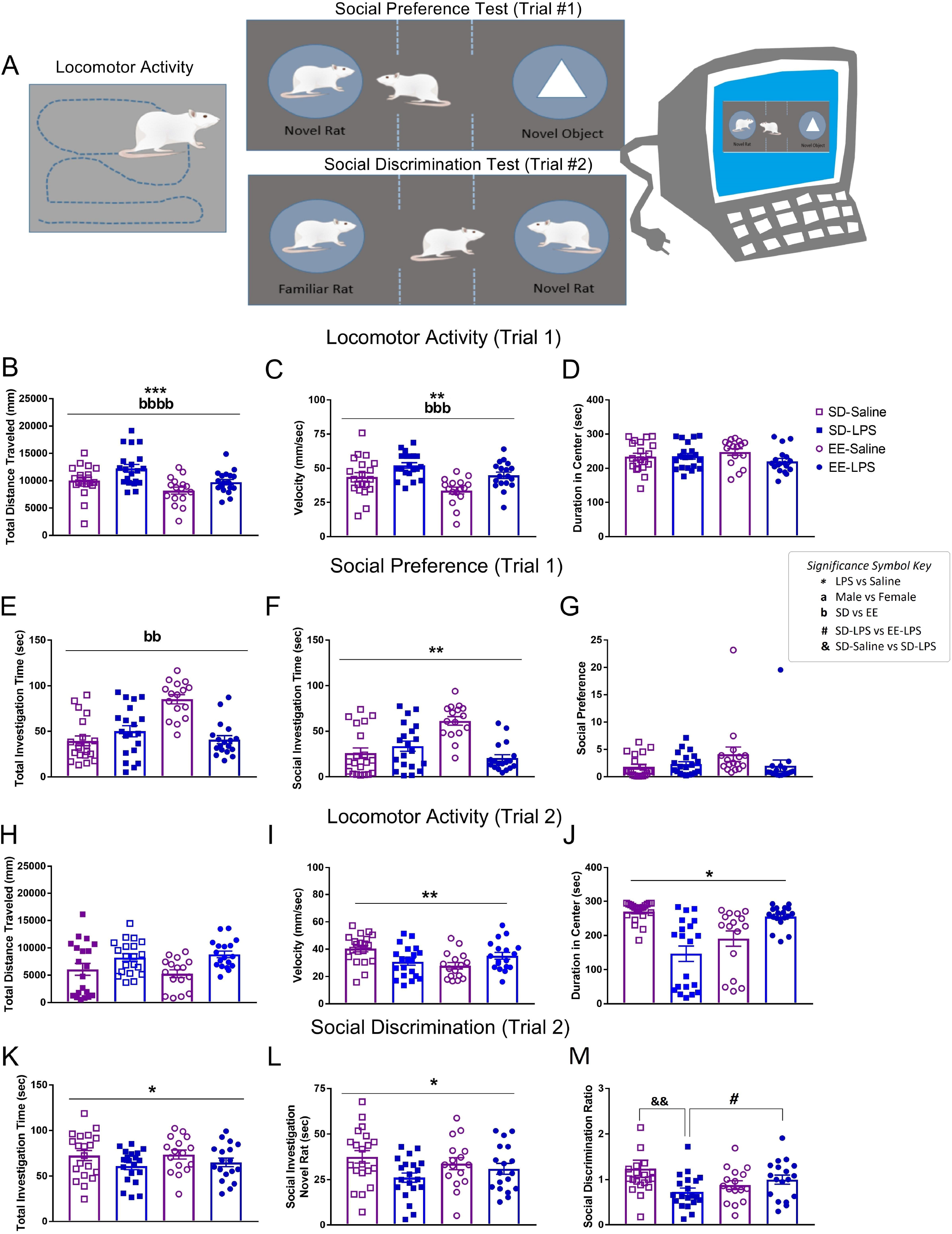
A) Illustration of juvenile locomotor activity, social preference (Trial 1), and social discrimination (Trial 2) tests using automated behavioral tracking software. B) Total distance traveled (mm), C) velocity (mm/sec), D) duration spent in center (sec), total time investigating (seconds) E) both the novel object and the novel conspecific, F) the novel conspecific, and G) the social preference score for Trial 1. H) Total distance traveled (mm), I) velocity (mm/sec), J) duration spent in center (sec), total time investigating (seconds) K) both the novel and familiar conspecifisc, L) the novel conspecific, and M) the social discrimination score for Trial 2. Group designations: SD-Saline (open squares), SD-LPS (closed squares), EE-Saline (open circles), EE-LPS (closed circles). Error bars represent mean ± SEM, n = 8-10 per sex/housing/MIA group. If there were no significant effects of sex, male and female offspring data were collapsed together for visualization. *p < 0.05, ** p < 0.01, *** p < 0.001, ****p <0.0001, LPS vs saline; ^a^p < 0.05, ^aa^p < 0.01, ^aaa^p < 0.001, ^aaaa^p <0.0001, male vs female; ^b^p < 0.05, ^bb^p < 0.01, ^bbb^p < 0.001, ^bbbb^p <0.0001, standard housing vs environmental enrichment; ^&&^p < 0.01, SD-Saline vs SD-LPS ^; #^p < 0.05, SD-LPS vs EE-LPS.

In the social preference test (Trial 1; **Figure 3A**), EE rats spent more time investigating (seconds spent with novel object + novel rat) than SD animals (*X*^2^(1) = 6.224, p =0.013; **Figure 3E**). In contrast, LPS animals (27.49 ± 3.53 sec) interacted less with social conspecifics compared to saline rats (41.66 ± 4.79 sec; *X*^2^(1) = 4.134, p =0.042; **Figure 3F**), although this did not translate into changes in social preference (p>0.05; **Figure 3G**).

While there were no effects on distance traveled in the social discrimination test (Trial 2; **Figure AH**), gestational LPS animals still traveled at faster speeds (F(1, 66) = 11.641, p = 0.001;; **Figure 3I**). During this trial, LPS animals spent less time in the center of the social chamber (*X*^2^(1) = 4.539, p = 0.033; **Figure 3J**) and, compared to saline animals, engaged in fewer investigative behaviors overall (*X*^2^(1) = 3.223, p =0.042; **Figure 3K**). There were no significant differences in the duration investigating the familiar rat (p >0.05; data not shown), however, LPS animals spent less time investigating the novel conspecific than their saline counterparts (F(1,66) = 5.249, p = 0.025;; **Figure 3L**). This translated into a significant gestational treatment by housing interaction on the social discrimination ratio (F(1,66) = 6.362, p = 0.014; **Figure 3M**). Social discrimination between the novel and familiar conspecifics was significantly impaired in SD-LPS compared to SD-Saline animals (t(38) = −2.849, p = 0.007). This gestational effect of LPS was not observed in EE housed animals (EE-LPS vs EE-Saline p > 0.05). Moreover, SD-LPS rats had lower discrimination ratios compared to EE-LPS (t(36) = −2.067, p = 0.046), suggestive of a protective effect of complex environments against MIA. Additionally, there were no sex differences, highlighting that female offspring are also vulnerable to social disruptions associated with MIA.

### MIA and enrichment effects on Crh mRNA in the juvenile brain

Although hippocampal *Crh* was unaffected by early life experience (p >0.05; **Figure 4A**), hypothalamic *Crh* mRNA was significantly higher in MIA animals compared to saline (*X*^2^(1) = 5.596, p = 0.018; **Figure 4B**).

**Figure 4.**
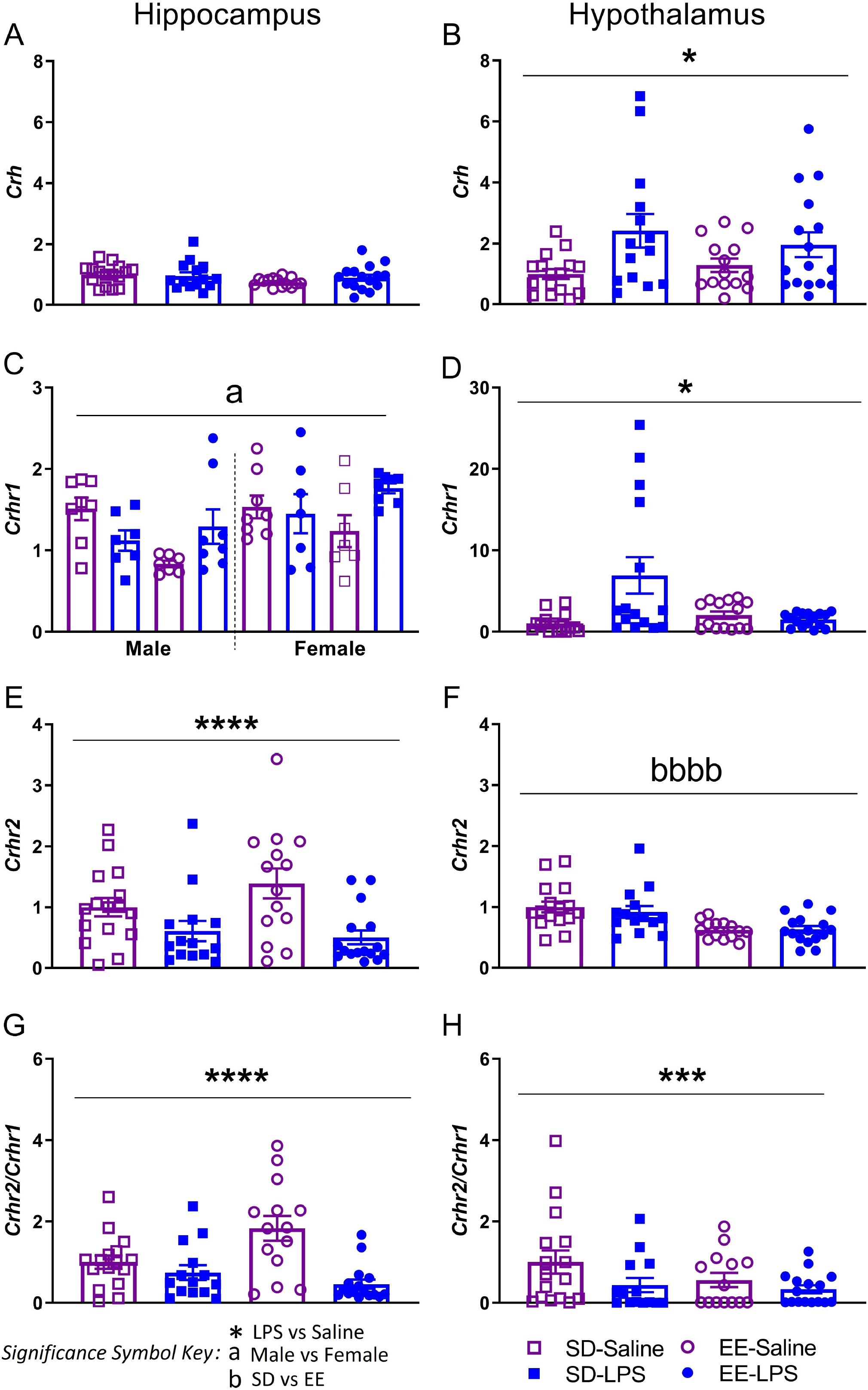
Juvenile brain mRNA expression of hippocampal (left) and hypothalamic (right) AB) *Crh*, CD) *Crhr1*, EF) *Crhr2*, GH) *Crhr2/Crhr1* normalized to *Gapdh*. Group designations: SD-Saline (open squares), SD-LPS (closed squares), EE-Saline (open circles), EE-LPS (closed circles). Error bars represent mean ± SEM, n = 7 per sex/housing/MIA group. If there were no significant effects of sex, male and female offspring data were collapsed together for visualization. *p < 0.05, ** p < 0.01, *** p < 0.001, ****p <0.0001, LPS vs saline; ^a^p < 0.05, ^aa^p < 0.01, ^aaa^p < 0.001, ^aaaa^p <0.0001, male vs female; ^b^p < 0.05, ^bb^p < 0.01, ^bbb^p < 0.001, ^bbbb^p <0.0001, standard housing vs environmental enrichment.

Females (1.51 ± 0.09) had higher levels of hippocampal *Crhr1* than males (1.21 ± 0.08; *X*^2^(1) = 6.466, p = 0.011; **Figure 4C**). Expression of *Crhr1* in the hypothalamus was elevated in LPS (10.53 ± 2.34) versus saline animals (4.89 ± 0.89; *X*^2^(1) = 4.103, p = 0.043; **Figure 4D**), but this effect was lost following application of the FDR correction. In contrast, hippocampal *Crhr2* expression was significantly reduced in MIA rats (*X*^2^(1) = 12.801, p =0.0001; **Figure 4E**). Surprisingly, *Crhr2* was elevated in SD animals (12.07 ± 0.98) compared to EE (8.26 ± 0.35; t(58) = 4.535, p = 0.0001; **Figure 4F**). Notably, the relative expression of hippocampal (*X*^2^(1)= 13.552, p = 0.0001; **Figure 4G**) and hypothalamic (X^2^(1) = 7.562, p = 0.006; **Figure 4H**) *Crhr2*/*Crhr1* was significantly reduced following MIA.

### MIA and enrichment effects on glucocorticoid and mu opioid receptors in the juvenile brain

Following the FDR correction, there were no significant differences in either hippocampal or hypothalamic *Nr3c1* (p > 0.05; **Figure 5AB**).

**Figure 5.**
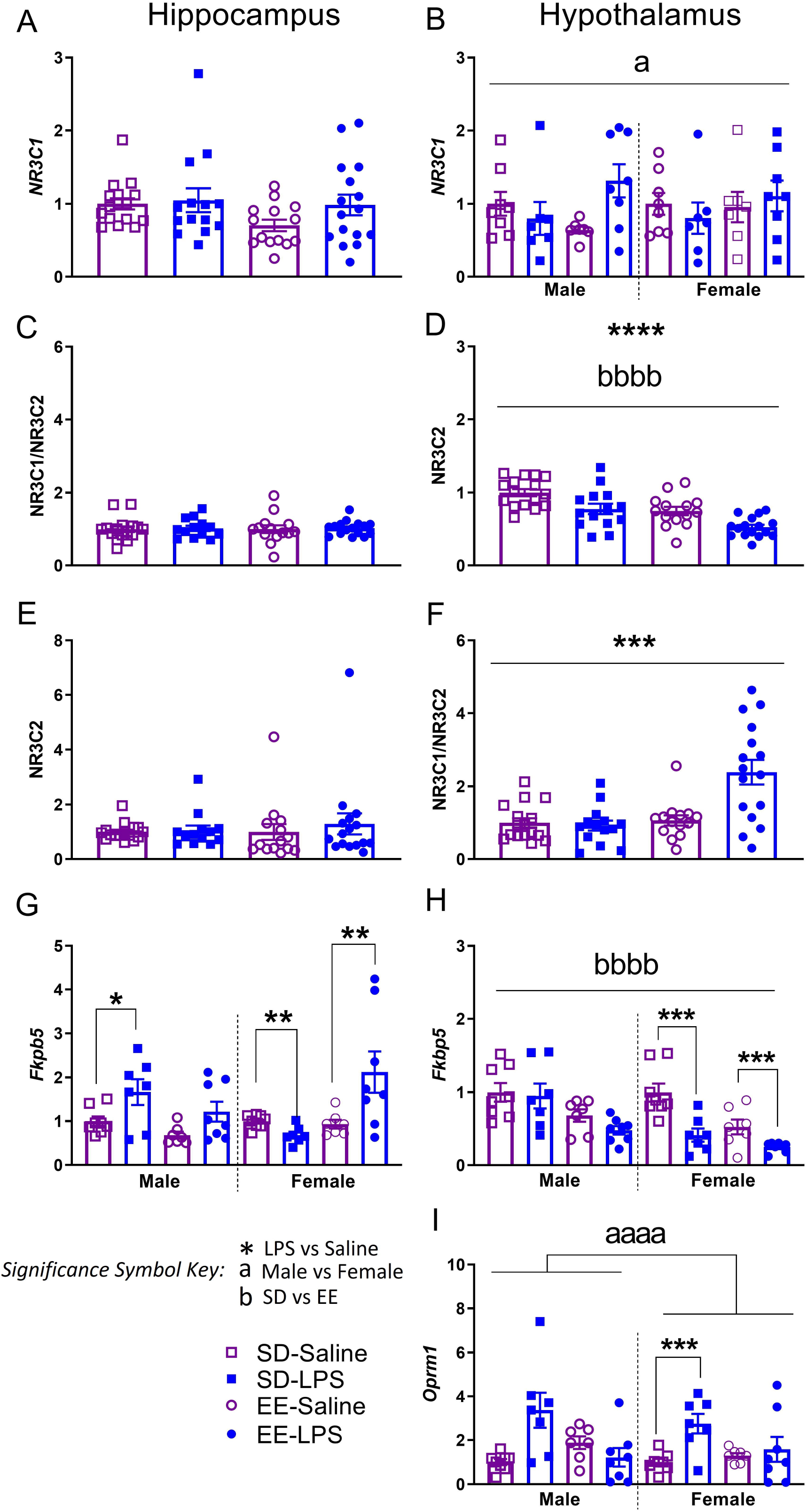
Juvenile brain mRNA expression of hippocampal (left) and hypothalamic (right) AB) *Nr3c1*, CD) *Nr3c2*, EF) *Nr3c1/Nr3c2*, GH) *Fkbp5*, and I) *Oprm1* normalized to *Gapdh*. Group designations: SD-Saline (open squares), SD-LPS (closed squares), EE-Saline (open circles), EE-LPS (closed circles). Error bars represent mean ± SEM, n = 7 per sex/housing/MIA group. If there were no significant effects of sex, male and female offspring data were collapsed together for visualization. *p < 0.05, ** p < 0.01, *** p < 0.001, ****p <0.0001, LPS vs saline; ^a^p < 0.05, ^aa^p < 0.01, ^aaa^p < 0.001, ^aaaa^p <0.0001, male vs female; ^b^p < 0.05, ^bb^p < 0.01, ^bbb^p < 0.001, ^bbbb^p <0.0001, standard housing vs environmental enrichment.

Hippocampal *Nr3c2, Nr3c1*/*Nr3c2*, and hypothalamic *Nr3c2* were not different across treatment groups (p > 0.05; **Figure 5CDE, *see Supplementary Table 3***). On the other hand, saline (t(58) = −3.911, p = 0.0001) and SD rats (t(58) = 4.209, p = 0.0001) had elevated hypothalamic expression of *Nr3c2* compared to LPS and EE animals respectively **(Figure 5D**). EE animals (0.47 ± 0.06) had significantly higher levels of *Nr3c1*/*Nr3c2* mRNA in the hypothalamus (*X*^2^(1) = 7.400, p =0.007; **Figure 5F**) compared to SD (0.26 ± 0.025); this effect seemed particularly driven by elevations of mRNA in EE-LPS animals, perhaps a compensatory effect against MIA challenge.

Male SD-LPS rats had elevated hippocampal *Fkbp5* compared male SD-Saline rats (t(13) = 2.253, p = 0.042; **Figure 5G**). This effect was not seen in similarly treated EE male animals (p >0.05), suggestive of environmentally mediated protection. In contrast to the males, female SD-LPS rats had lower hippocampal *Fkbp5* levels compared to female SD-Saline animals (t(13) = −3.565, p = 0.003). A similar effect was seen with hypothalalmic *Fkbp5* where MIA females had lower expression compared to saline (t(28) = −4.108, p = 0.001; **Figure 5H**), regardless of housing condition. This may indicate that female rats are more resilient to stress-induced disruptions to the hypothalamic glucocorticoid system.

Interestingly, female EE-LPS rats had heightened levels of *Fkpb5* in the hippocampus compared to EE-Saline females (t(13) = −3.562, p = 0.005; **Figure 5G**); perhaps EE is not as ‘enriching’ for females, or acts as an aversive stimulus in a brain circuit-dependent manner, potentiating the detrimental effects of MIA on some measures. That said, hypothalamic Fkbp5 was significantly elevated in male and female SD rats (0.11 ± 0.01), compared to EE (0.06 ± 0.01; t(58) = 4.023, p = 0.0001; **Figure 5H**), underscoring the possibility that standard laboratory housing conditions may be inherently stressful.

SD-LPS animals had elevated levels of hypothalamic *Oprm1* compared to SD-Saline rats (t(28) = 3.189, p = 0.004; **Figure 5I**). EE-Saline and EE-LPS animals were not significantly different (p >0.05), suggestive of a modestly protective effect of this rearing condition on *Oprm1* expression following MIA.

Finally, female LPS (3.55 ± 0.65; t(28) = −3.967, p = 0.001) and female saline (1.9 ± 0.17; t(28) = −7.421, p = 0.0001) treated animals had higher hypothalamic *Oprm1* compared to male LPS (0.86 ± 0.20) and male saline animals (0.55 ± 0.07), respectively (**Figure 5I**).

### MIA and enrichment effects on epigenetic writers in the juvenile brain

There were no significant effects of early life experience on hippocampal *OGT*, *DNMT1*, *DNMT3a*, or *MECP2* expression (p>0.05; **Table 1**). However, hypothalamic *OGT* (*X*^2^(1) = 7.122, p =0.008; **Table 1**), *DNMT1* (*X*^2^(1) = 19.478, p = 0.001; **Table 1**), and *DNMT3a* (*X*^2^(1) = 20.136, p =0.0001; **Table 1**) expression was lower in MIA exposed animals compared to saline. A three-way interaction was revealed for hypothalamic *MECP2* (F(1, 52) = 22.305, p =0.0001;). Females treated with LPS had lower *MECP2* than female saline rats (SD animals: t(13) = −2.577, p =0.023; EE animals: t(13) = −4.5724, p = 0.002; **Table 1**). Additionally, female EE rats had higher hypothalamic *MECP2* levels than both male EE (t(12) = −4.754, p = 0.0001; **Table 1**) and female SD rats (t(13) = −5.368, p =0.0001; **Table 1**).

## Discussion

This study provides the first evidence that EE can impact placental functioning, potentially protecting the developing offspring from the consequences of stress-induced fetal programming. While there is a plethora of data demonstrating that gestational stressors affect placenta and fetal development (Brunton & Russell, 2011; Chen & Gur, 2019; Monk et al., 2012; St. Pierre et al., 2018), a stronger emphasis on identifying interventions to promote maternal/offspring protection from adverse experiences is needed (Kentner et al., 2019b; Shonkoff et al., 2000).

As expected (Asiaei et al., 2011; Kirsten et al., 2013), maternal plasma corticosterone and IL-1β were elevated 3 h after LPS challenge on gestational day 15. We did not measure maternal IL-6 levels as this proinflammatory cytokine is not consistently induced by LPS during the early-to-mid gestational period, even in the presence of MIA-associated behavioral and neurophysiological disruptions in offspring (Asiaei et al., 2011; Elovitz et al., 2011). Evidence for MIA-associated glucocortocoid programming (Seckl & Meaney, 2004; Heussner et al., 2016) was demonstrated by a simultaneous decrease of both *Hsd11b2* and *Hsd11b2/Hsd11b1* in our male and female placentas. These placental, but not maternal, implications of MIA were attenuated by EE. Moreover, we observed changes in central *Fkbp5* expression in juvenile MIA offspring, particularly in male hippocampus which showed a phenoytype suggestive of altered stress responsivity (Binder, 2009; Matosin et al., 2018). This effect was also ameliorated by complex housing.

Given that life-long EE exposure attenuates the MIA-induced downregulation of glucocorticoid receptor protein in juvenile male hippocampus (Connors et al., 2014), we anticipated a similar effect with *Nr3c1* in young juvenile animals. While we observed no indication of *Nr3c1* downregulation following MIA, associated changes in hippocampal *Fkbp5* are suggestive of significant alterations in glucocorticoid receptor sensitivity (Binder, 2009). Combined with disrupted hypothalamic *Nr3c2* and *Nr3c1/Nr3c2* levels, it is clear that there are circuit-dependent consequences of early life inflammatory exposure. Moreover, these consequences (both behavioral and neurophysiological) emerge across differential trajectories via interactions with developmental variables such as puberty (Clark et al., 2019; Dinel et al., 2014; MacRae et al., 2015; Zuckerman et al., 2003; Meyer et al., 2006b). Enrichment also interacts with these developmental periods, as evidenced by elevated *Nr3c1* in fetal, but not prepubertal EE juvenile brain. Therefore, it may be that the downregulation and rescue of *Nr3c1* following MIA is only observable later in adolescent development. Alternatively, while 11HSD2 activity is believed to parallel the pattern of mRNA expression (Diaz et al., 1998), it is important to note that mRNA is not always reflective of protein amounts (Chen et al., 2002; Greenbaum et al., 2003), and the *Nr3c1* gene itself may be unaffected by prenatal LPS challenge.

While EE appeared to have a discrete influence on glucocorticoid-associated placental mediators such as *Hsd11b2*, this intervention may have instead exerted its protective influence through separate molecular or immune pathways. For example, placental hemorrhage, hypoxia, interruptions in neural progenitor cell proliferation, tumor necrosis factor-α, and the maternal-fetal leukemia inhibitory factor (LIF) signal pathway have all been implicated in LPS-induced disruptions of fetal brain development (Izvolskaia et al., 2018; Tsukada et al., 2019). Moreover, in poly I:C MIA models, IL-6 (Smith et al., 2007; Hsiao & Patterson, 2011) and IL-17a/activated Th17 cells (Choi et al., 2016) are well recognized to underlie the associated phenotypes in offspring, as are changes in placental morphology (Murray et al., 2019). Of course, this does not preclude the notion that EE may have a combined effect on multiple or even redundant pathways, leading to protection of the placenta and the fetus. The protective potential of EE on placental functioning including tight junctions, glucose transport capacity and hormone secretion following MIA remains to be tested.

The therapeutic benefits of EE may also be dependent on the immunogen challenge. For example, previous work using polyriboinosinic polyribocytidylic acid (poly I:C) suggests that MIA may counteract the advantages of complex housing (Buschert et al., 2016). However, inconsistent findings between our two studies may instead be due to methodological differences such as timing of first EE exposure (e.g. preconception vs post weaning) and species employed. While we use rats, Buschert and colleagues (2016) employed male CD-1 mice which are well known to become aggressive when housed in EE (McQuaid et al., 2012; McQuaid et al., 2013). The stress associated with territorial aggression (e.g. fighting for resources/EE devices) and unstable social hierarchies may account for the blocked action of EE in response to MIA. Future studies will need to better determine the generalization of EE across immunological challenges, species, and exposure timing.

As mentioned, the benefits of EE may also be dependent on the critical period of exposure in addition to interactions with other variables such as parental care. Indeed, EE during isolated developmental periods (e.g. pre-conceptive/reproductive, prenatal, preweaning) is known to affect the early nurturing behaviors of parents and subsequent offspring outcomes (Cutuli et al., 2018). For example, parental EE is associated with observed effects on nursing quality and quantity, maternal contact time with the nest (Connors et al., 2015; Curley et al., 2009; Korgan et al., 2016; Welberg et al., 2006; Rosenfeld and Weller, 2012; Zuena et al., 2016), in addition to reports of changes in licking and grooming (Cancedda et al., 2004; Sale et al., 2004). In turn, progeny demonstrate accelerated motor (Caporali et al., 2014) and visual (Cancedda et al., 2004; Sale et al., 2004) development, in addition to sex-specific alterations in learning, memory, anxiety, and social behaviors (Connors et al., 2015; Rosenfeld and Weller, 2012; Welberg et al., 2006; Zuena et al., 2016). These findings highlight the difficulty in dissociating the effects of EE from parental care to test if enrichment exposure during a particular developmental period is sufficient to offset the long-term effects of MIA in offspring. This task is challenged further by data showing that EE removal (and replacement into SD conditions) may simulate the experience of loss, inducing a depressive-like pheonotype (Smith et al., 2017; Morano et al., 2019). When EE is removed from dams, this may increase maternal care as a form of stimulation replacement (Rosenfeld and Weller 2012; Cutuli et al., 2018). Even in humans, EE interventions (Woo & Leon, 2013; Woo et al., 2015; Aronoff et al., 2016 Morgan et al., 2013; Morgan et al., 2015; Purpura et al., 2014) may be impacting parental-offspring attachments, underlying their benefits (Kentner et al., 2019b). Thus, future laboratory work will require cross-fostering animals from their dams to explore these questions further.

Epigenetic influences likely underlie the observed fetal protection from MIA as they are known to be affected by both maternal care (Zhang et al., 2013; Stolzenberg & Mayer, 2019; Miguel et al., 2019) and EE (Zhang et al, 2018). Moreover, in the placenta, prenatal stress increased transcription of DNMTs and CpG methylation of the *Hsd11b2* gene (Jensen Peña et al., 2012). Both 3 and 24 h following MIA, we observed elevated levels of placental *DNMT1* (independent of inflammation) in EE exposed animals, suggestive of enduring experience-dependent effects on placental functioning. Twenty four hours after LPS challenge, placental enzymes were mostly normalized, except for a modest elevation in *Hsd11b2* observed in EE placentas. To our knowledge, this is the first evidence of EE protecting the placenta directly, which may influence later offspring behavior as the placenta is recognized to regulate the prenatal environment and infant outcomes (Lesseur et al., 2014).

Elevations in placental *MECP2* did not become apparent until 24 h following inflammatory challenge, highlighting the time-dependent nature of epigenetic changes. Dysregulation of this chromatin-associated MeCP2 protein can lead to elevations in DNA methylation of the *glut3* gene and interruptions in trans-placental glucose transport, affecting fetal development (Ganguly et al., 2014). Offspring of mouse dams treated with poly I:C presented with both hyperW- and hypomethylated CpGs at several loci and at at distinct genomic regions (Richetto et al., 2017). These methylation changes corresponded to transcriptional changes of genes relevant to GABAergic differentiation and signaling, Wnt signaling, and neural development. Notably, the expression of these effects was dependent on the critical period of poly I:C challenge (mid vs late pregnancy) and age of the offspring.

We also observed a sex-specific effect of lower *OGT*, a marker of cellular stress during pregnancy, in male compared to female placenta; an effect reported previously (Howerton et al., 2013; Howerton & Bale, 2014). The sexual dimorphism of placental OGT expression appears to be influenced by the type of maternal stressor and its magnitude (Lima et al., 2018). Most interestingly, we found *OGT* to be elevated in female placentas 24 h after exposure to MIA, suggestive of a compensatory protective mechanism which could account for reported female resilience in MIA models. A similar pattern of putative protection was observed for juvenile hypothalamic *Fkbp5*, which was lower in females exposed to MIA. Importantly, some of the foundational information reported here lays the groundwork for future larger epigenetic studies (e.g. bisulfite sequencing to evaluate methylation patterns of *Fkbp5, Nr3c1, Crh, Opmr1*) as the significant changes in juvenile mRNA expression could be due to epigenetic modification of these genes. Further work is required to determine if the long-term influence of EE on offspring behavior is mediated via broad changes in DNA methylation and protein glycosolation in placenta.

Although males are thought to be more vulnerable to MIA (Meyer, 2014; Reisinger et al., 2015), and despite evidence of epigenetically conferred protection to our female animals, we demonstrate that both sexes can be affected by prental LPS challenge. Here, both male and female offspring had significant impairments in social discrimination ability. These data contribute to a growing literature suggesting that female animals are vulnerable to early life insults, which may become apparent in a test- or age-specific manner. For example, using a poly I:C MIA model (Patrich et al., 2016), juvenile male rats showed depression in hippocampal excitatory transmission, an effect that did not appear in poly I:C exposed females until adulthood. Moreover, while male (but not female) exposed poly I:C offspring displayed disrupted startle responses to acoustic stimuli, both sexes showed impairments in sociability, visual discrimination and novel object preferences (Lins et al., 2018; Lins et al., 2019). Reported sex differences are likely due to timing of infection, since male and female brains do not develop at the same pace (Anderson, 2003). In general, MIA associated behavioral and neurophysiological outcomes may appear differentially, or similar behavioral phenotypes can be mediated by separate underlying mechanisms (Sorge et al., 2015), as is probable with the social discrimination impairments observed here. Notably, the paucity of female impairments reported following MIA are likely due to experimental biases, leading to the exclusion of female animals in research (Coiro & Pollak, 2019) and deficits in the detection of such impairments.

Going forward it will be necessary to to identify the specific elements of EE (e.g. exercise, novelty, social activity) that impact the dam, placenta and/or fetus to confer protection. This will help with the development of targeted interventions that may generalize clinically (Kentner et al., 2018b; Kentner et al., 2019b). Combined with evidence that EE attenuated the placental mediated programming effects of the glucocorticoid system following MIA, these data provide the first report that enrichment protects placental functioning at the time of a maternal stressor. Future work will need to evaluate the optimal periods of EE use, to determine if enrichment exposure during pregnancy is sufficient, or if there is an interaction with postnatal experience. Overall, changes in placenta functioning offer a putative mechanism by which the positive care of mothers can lead to a beneficial or protective outcome for the offspring. This underscores the importance of supporting maternal health throughout pregnancy and beyond.

## Acknowledgments

The authors would like to extend their thanks to Antoine Khoury and Molly MacRae for technical assistance.

