## Supplementary Table 2 for "Environmental influences on placental programming and offspring outcomes following maternal immune activation"

**Supplementary Table 2.** Fetal brain mRNA gene expression

| **Sex** | **Housing** | **Gestational Treatment** | **3 h post LPS** | | **24 h post LPS** | |
| --- | --- | --- | --- | --- | --- | --- |
|  |  |  | ***NR3C1*** | ***NR3C2*** | ***NR3C1*** | ***NR3C2*** |
| *Male* | *SD* | *Saline* | 1.00±0.41 | 1.00±0.56 | 1.00±0.82 | 1.00±0.688 |
|  |  | *LPS* | 0.02±0.01 | 0.05±0.02 | 4.43±1.75 | 1.48±0.56 |
|  | *EE* | *Saline* | 0.50±0.31 | 1.34±0.79 | 2.70±1.30*** | 2.81±1.17*** |
|  |  | *LPS* | 30.41±14.94 | 15.10±7.55 | 3.27±1.30*** | 5.14±1.16*** |
| *Female* | *SD* | *Saline* | 1.00±0.74 | 1.00±0.23 | 1.00±0.16 | 1.00±0.35 |
|  |  | *LPS* | 1.42±1.01 | 6.30±4.38 | 3.34±0.84 | 2.05±0.76 |
|  | *EE* | *Saline* | 14.71±9.83 | 3.44±1.91 | 26.92±9.29*** | 8.63±2.88*** |
|  |  | *LPS* | 16.46±10.70 | 4.04±3.04 | 15.73±7.03*** | 7.46±2.43*** |

Error bars represent mean ± SEM, n = 7 per group. ^*^p < 0.05, ^**^p < 0.01, ^***^p < 0.001, ^****^p <0.0001 significantly different from SD. Data normalized to same-sex SD-Saline groups.
