## Supplementary Table 3 for "Environmental influences on placental programming and offspring outcomes following maternal immune activation"

Supplementary Table 3. Extra Omnibus Reporting Statistics

| **Variable** | **Effect** | **Statistics** |
| --- | --- | --- |
| Distance Traveled – Trial 1 | Gestational Treatment | *X*^2^(1) = 4.820, p = 0.028 |
|  | Housing | *X*^2^(1) = 10.801, p = 0.001 |
| Velocity – Trial 1 | Gestational Treatment | F(1,66) = 13.859, p = 0.001 |
|  | Housing | F(1,66) = 10.580, p = 0.002, |
| Social Preference – Trial 1 | Sex x Gestational Treatment | F(1, 66) = 4.851, p = 0.031; follow up tests N.S |
| Hypothalamic *Crhr2* | Housing | F(1, 52) = 20.611, p = 0.0001 |
| Hippocampal *Nr3c2* | Sex | F(1, 52) = 4.453, p = 0.040; follow up tests N.S |
| Hypothalamic *Nr3c2* | Gestational Treatment | F(1, 52) = 19.049, p = 0.0001 |
|  | Housing | F(1, 52) = 21.702, p = 0.0001 |
| Hippocampal *Nr3c1*/*Nr3c2* | Sex x Gestational Treatment | F(1, 51) = 4.526, p = 0.038; follow up tests N.S |
| Hippocampal *Fkbp5* | Sex x Gestational Treatment x Housing | F(1, 52) = 5.441, p = 0.024 |
| Hypothalamic *Fkbp5* | Sex x Gestational Treatment | F(1, 52) = 10.707, p = 0.002 |
| Hypothalamic *Oprm1* | Gestational Treatment x Housing | F(1, 52) = 7.455, p = 0.009 |
|  | Sex x Gestational Treatment | F(1, 52) = 4.251, p = 0.044 |
